# Commercially Purchased and In-House Bred C57BL/6 Mice with Different Gut Microbiota Exhibit Distinct Indomethacin-Induced Toxicities

**DOI:** 10.1101/2025.05.01.651704

**Authors:** Jianan Zhang, Rose Viguna Thomas Backet, Josh J. Sekela, Meredith J. Zeller, Rani S. Sellers, Matthew R. Redinbo, Ajay S. Gulati, Aadra P. Bhatt

## Abstract

Non-steroidal anti-inflammatory drug (NSAID)-induced toxicities are a significant clinical problem, yet the factors influencing these outcomes remain incompletely understood. Here, we investigated the impact of mouse vendor on indomethacin-induced injury using C57BL/6 mice from different breeding facilities (in-house “Tar Heel” and commercial Charles River). We found that Tar Heel mice exhibited significantly enhanced susceptibility to indomethacin toxicity, characterized by greater body weight loss, increased ileal ulceration, elevated fecal lipocalin-2 levels, and higher goblet cell numbers in ileum compared to Charles River mice.

Importantly, whole genome metagenomic analysis revealed distinct baseline gut microbiomes between the two types of mice. Notably, Tar Heel mice showed higher abundances of β-glucuronidase (GUS)-producing bacteria, particularly those expressing Loop-1 GUS enzymes, and elevated levels of mucolytic enzyme-encoding bacteria. These differences suggest that enhanced indomethacin toxicity observed in Tar Heel mice may be related to functional changes in their gut microbiome, which may predispose to an exaggerated response to NSAID exposure. Together, our findings demonstrate that vendor-specific differences significantly influence NSAID-induced intestinal toxicity and highlight the importance of considering mouse sources and gut microbial compositions in experimental design. Moreover, we highlight potential functional roles that gut microbes play in host-indomethacin interactions.

## INTRODUCTION

Gastrointestinal (GI) toxicity is a significant side effect of non-steroidal anti-inflammatory drugs (NSAIDs), such as indomethacin, which are utilized by millions of patients worldwide. As many as 16,500 deaths annually in the US are caused by NSAID-induced GI bleeding ^1-3^. Beyond the well described gastric damage, NSAIDs also cause enteropathy resulting from the interactions between host physiology and the gut microbiome, as well as the biochemical properties of the NSAIDs themselves ^4-6^. NSAIDs are amphiphilic molecules whose physicochemical properties allow them to penetrate through both the intestinal mucus layer and phospholipid-rich epithelial cell membranes. NSAIDs can result in increased mucosal exposure to luminal aggressors such as gut bacteria. At physiologically relevant concentrations, NSAIDs are known to cause mitochondrial uncoupling, leading to increased death of intestinal stem- and differentiated cells, and increased intestinal permeability ^7,8^.

Gut microbiota have an additional role in NSAID-induced toxicity. A range of NSAIDs, including indomethacin, are conjugated with glucuronic acid in the liver and excreted into the intestine as inactive glucuronide metabolites. Many intestinal bacteria species produce β-glucuronidase (GUS) enzymes that deconjugate these metabolites, thereby reactivating the drug and increasing local gut luminal concentrations, which further contribute to intestinal damage ^9^. The abundance and activity of GUS-producing bacteria in the GI tract can therefore significantly impact the severity of NSAID-induced intestinal damage ^10,11^. Furthermore, host factors such as differential COX activities, responses to redox stress, innate immunity, and the induction of inflammatory cytokines lead to further epithelial erosions and ulceration ^12^.

Gut microbiota functionally influence the toxicity profile of NSAIDs in the host. Intestinal mucus, primarily produced by goblet cells, forms a crucial protective barrier against luminal contents and drug-induced injury. Some bacterial species produce mucolytic enzymes that can degrade this protective layer ^13^. In the context of NSAID use, an overabundance of these mucolytic bacteria may compromise the mucus barrier, exacerbating drug-induced damage to the underlying epithelium. Moreover, these microbial factors interact with the host, particularly with structures like goblet cells, which are essential for maintaining intestinal homeostasis ^14^. NSAID-induced alterations in goblet cell function or number^15^, potentially influenced by the local microbial environment, could further contribute to gut toxicity.

Animal models, particularly mice, are invaluable tools for investigating NSAID-induced GI toxicity and potential therapeutic interventions ^16,17^. These models facilitate the study of mechanisms involved in drug-induced damage and can test novel protective strategies under controlled conditions. However, the source of laboratory animals—whether they are bred in-house or obtained from commercial suppliers—may introduce variability in experimental outcomes. Advances in metagenomic sequencing technologies have revolutionized our ability to characterize complex microbial communities with unprecedented depth and resolution. Whereas previous studies showed that differential microbiota composition between vendors is an important consideration in the rodent studies ^18,19^, the mechanisms driving these differences remain unclear.

Understanding the interplay between the function of the gut microbiome, their enzymatic activities, and host cellular responses is crucial for elucidating the mechanisms of NSAID-induced GI toxicity, with the goal of preventing or mitigating this damage. Moreover, differences in the abundance or activity of these key microbial players between in-house bred and commercially sourced mice could significantly impact experimental outcomes in NSAID toxicity studies. In this study, we examine differences in the susceptibility to indomethacin-induced injury of in-house-bred versus commercially purchased mice. We further investigate how microbial community structure and function differ between these groups. Our findings advance the mechanistic understanding of interactions between NSAIDs, the gut microbiome, and host physiology. These results could benefit the reproducibility and translatability of preclinical studies using different mouse sources and shed light on the possible functional role of specific microbial species in modulating NSAID toxicity.

## MATERIALS & METHODS

### Animals

All animal experiments were conducted in accordance with the protocols approved by the Institutional Animal Care and Use Committee of the University of North Carolina at Chapel Hill (IACUC approval numbers: 22-170.0 and 21-049.0). Mice maintained in a specific pathogen free (SPF) facility with 12-hour light/dark cycle, maintained between 20-23°C with 30-70% relative humidity. Mice had ad libitum access to drinking water and chow (irradiated Purina PicoLab® Select Rodent 50 IF/6F 5V5R*) for the entire study. In-house bred male and female “Tar Heel” mice were bred in the AAALAC-approved University of North Carolina at Chapel Hill (UNC) animal facility and were originally purchased from the Jackson Laboratory (C57BL/6J). Charles River (C67BL/6N) male and female mice were commercially bred and purchased from Charles River Laboratories. Externally purchased mice were allowed to acclimatize for two weeks in our UNC animal facility prior to further experiments.

### Animal Experiment 1: Effects of indomethacin on a mouse model in ileum

Tar Heel mice (male, 8-10-week-old) were obtained from the UNC facility and allowed to acclimatize for two weeks prior to study initiation. Mice were randomly assigned into two groups: those orally gavaged with indomethacin (dose = 10 mg/kg body weight; n=8) or gavaged with vehicle (Veh; n=7). Gavages were performed once daily in the morning for 3 days (experimental design in **Fig 1A**). Vehicle was a sterile solution consisting of 10% dimethylformamide, 0.5% methyl cellulose and 0.5% tween-80. Mice were euthanized 24 hours following their last dose of indomethacin or vehicle, and blood and tissues were collected and stored at -80°C for further analysis. Fecal samples were collected daily and immediately placed on dry ice prior to storage at -80°C for further analysis.

**Fig. 1.**
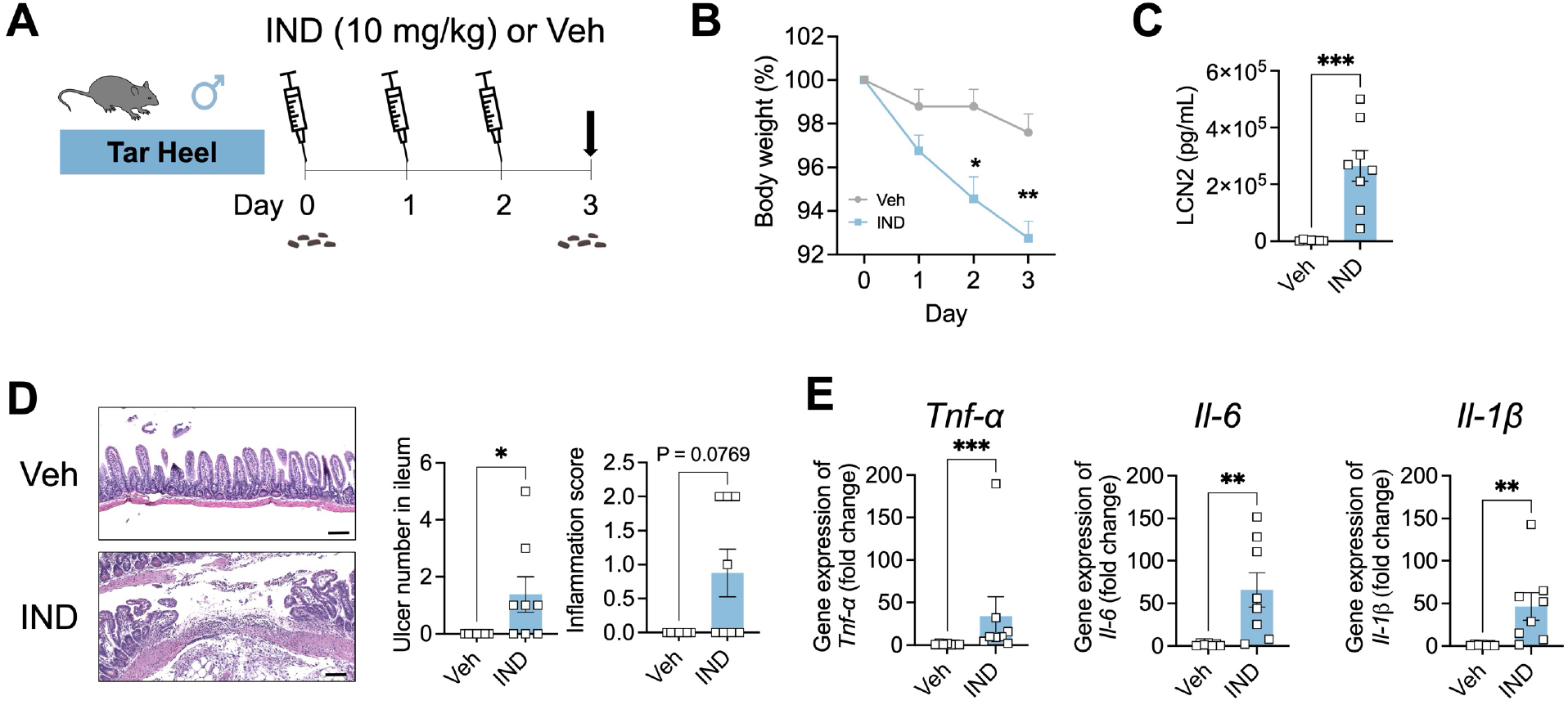
Three-day indomethacin treatment produces gastrointestinal toxicity. **A**. Scheme of experimental design in which in-house bred “Tar Heel” C57BL/6 male mice were treated with indomethacin (IND; 10 mg/kg body weight) or vehicle for 3 days, then euthanized 24 h after the last dose (dark arrow). Fecal material were collected at day 0, as indicated. **B**. Percent of day 0 body weight shows significant weight loss in IND treated mice at days 2 and 3. **C**. Fecal lipocalin-2 (LCN2) levels at day 3 are significantly higher in IND treated mice. **D**. Ulcers were found in ileum of IND treated mice but not vehicle treated mice. Representative H&E histological images of ileum are shown at left (scale bar = 100 μm), while at right the ulcer numbers in the ileum are significantly higher in IND treated mice and are absent in vehicle treated animals. **E**. Gene expression of pro-inflammatory cytokines *Tnf-α, Il-6*, and *Il-1β* normalized by *β-actin* in the ileum are significantly higher in IND treated mice than vehicle treated animals. n=7 in Veh group; n=8 in IND group. For the comparison between treatment groups, statistical significance was determined using two-side t-test. *P<0.05, **P<0.01, ***P<0.001, ****P<0.0001.

### Animal Experiment 2: Effects of indomethacin on in-house bred and commercial bred mice in ileum

Tar Heel and Charles River mice (male and female, 8-10-week-old) were obtained and allowed to acclimatize for two weeks prior to study initiation. Tar Heel and Charles River mice were randomly assigned into two groups for both sexes (n=11-19/group/sex): those orally gavaged with indomethacin (10 mg/kg body weight) or vehicle (Veh) for 3 total days as described in

**Animal Experiment 1** (experimental design in **Fig S1** and **2A**). Mice were euthanized 24 hours following their last dose of indomethacin or vehicle, and blood and tissues were collected and stored at -80°C for further analysis. Fecal samples were collected daily and immediately placed on dry ice prior to storage at -80°C for further analysis.

### Animal Experiment 3: Comparative metagenomic analysis of gut microbiomes on in-house bred and commercial bred mice

Tar Heel and Charles River mice (male and female, 8-10-week-old) were obtained and acclimatized as above. To investigate the microbial basis for differential indomethacin responses, we performed baseline gut microbiome analysis prior to drug treatment (experimental design in **Fig 4A**). Fresh fecal samples were collected from individual mice, flash-frozen on dry ice, and stored at -80°C until metagenomic sequencing analysis.

### Measurement of Lipocalin 2 (LCN-2) in Feces

Fecal samples were thawed, weighed and weight-normalized with PBS containing 0.1% Tween 20, then vortexed for 10 to 15 seconds. Samples were placed on a shaker in a cold room overnight, and the supernatant was collected after centrifugation at 12,000 rpm for 10 min. The content of LCN-2 in the supernatant was determined using a Mouse Lipocalin-2/NGAL DuoSet ELISA kit (R&D Systems, Catalog#DY1857) according to the manufacturer’s instructions. The optical density was determined using a microplate reader (CLARIOstar Plus) at 450 nm and 540 nm. The wavelength correction was determined by subtracting readings at 540 nm from the readings of 450 nm. LCN-2 concentration was calculated using a standard curve and expressed as pg/ml per mg of fecal material.

### Reverse-Transcriptase-qPCR of Inflammatory Biomarkers in Ileum

Two centimeters of ileal tissue was obtained from each mouse and frozen at -80 °C with RNAlater (Invitrogen). Total RNA was isolated using RNeasy Mini Kit (Qiagen) according to the manufacturer’s instructions. RNA was reverse transcribed into cDNA using SuperScript III reverse transcriptase (Invitrogen). 20 ul PCR reactions were prepared using Applied Biosystems SYBR green Master Mix (Thermo Fisher Scientific), and qPCR was carried out using a DNA Engine Quantstudio 3 system (Applied Biosystems). The sequences of mouse-specific primers (Sigma Aldrich) used to detect inflammatory biomarkers are listed in **Table S1**. The results for the target genes were normalized to *β-actin* using the 2 ^-ΔΔCT^ method.

### H&E Histological Staining and Ulceration Measurement in Ileum

The ileum of each mouse was opened longitudinally, Swiss rolled, and fixed in 10% phosphate buffered formalin (Thermo Fisher Scientific) for 48 hours. Tissue sections were routinely processed to paraffin, sectioned at 5 μm thickness every 100 μm for 3 times between sections, and stained with stained with hematoxylin and eosin (Epredia Richard-Allan Scientific). Samples were evaluated microscopically (RSS). Samples were assessed using a qualitative scoring system, except for mucosal erosions/ulcerations which were counted to give a total number in the sample. All three swiss roll samples were evaluated, and mucosal erosions/ulcerations were counted but not duplicated across sections (i.e. 3 sections have a lesion in the same location, lesion only counted once). Samples were scored according to **Table 1**.

**Table 1:** Criteria used to score microscopic findings in the ileum

| Finding | 1 | 2 | 3 | 4 | 5 |
| --- | --- | --- | --- | --- | --- |
| Inflammation (not including regions of ulceration) | Minimal and limited to LP | Mild and limited to LP or minimal but extending into SM | Mild in LP and SM | Moderate to severe in LP & SM or mild + crypt abscesses | Transmural |
| Distribution of most severe finding | <5%* | 6-30% | 31-60% | >60% | NA |
| Crypt degeneration and atrophy | minimal | mild | moderate | marked | severe |
| Distribution of crypt degeneration and atrophy | <5% | 6-30% | 31-60% | >60% | NA |
| LP = lamina propria of the ileum<br>SM = submucosa of the ileum<br>NA = not applicable<br>* Percent is an estimate |  |  |  |  |  |

### PAS-Alcian Blue Staining and Goblet Cell Quantification

Ileal tissue blocks were prepared from 10% phosphate buffered formalin fixed tissues, sectioned (5 μm) and stained with PAS-Alcian blue. Slides were examined and imaged under a light microscope using cellSens software (Olympus). Using ImageJ, goblet cells in the ileum were manually quantified for both number and area by analyzing at least 10 crypt-villus units per mouse.

### DNA Extraction from Fecal Samples

DNA was extracted from mouse fecal samples using the QIAmp DNA Stool Mini Kit (Qiagen, Valencia, CA) following instructions from the manufacturer with an additional bead-beating step ^20-22^. The quantity of the extracted DNA was measured using a NanoDrop Spectrophotometer (Thermo Fisher Scientific), and the quality was assessed on 1% agarose gels.

### Whole-Genome Sequencing and Analysis

PCR products were detected on 2% agarose gels by electrophoresis and purified using the Qiagen Gel Extraction Kit (Qiagen, Germany). Sequencing libraries were generated using NEBNext Ultra DNA Library Pre-Kit for Illumina, following manufacturer’s recommendations, and index codes were added. The library quality was assessed using the Qubit 2.0 Fluorometer (Thermo Fisher Scientific) and the Agilent Bioanalyzer 2100 system. The library was sequenced on an Illumina platform, and whole-genome reads were generated (10 Gb of raw data).

Metagenomic data were processed using MetaPhlAn (v4.06) to create a merged taxonomic relative abundance table ^23^. Alpha diversity was calculated by the Shannon diversity index using MetaPhlAn, and the Kruskal-Wallis rank sum test was employed to compare alpha diversity between groups. Beta diversity was calculated by the Bray-Curtis dissimilarity matrix using MetaPhlAn, and PERMANOVA testing was applied to compare beta diversity between groups using Adonis2. Classical multidimensional scaling was applied to the dissimilarity matrix to create a PCoA plot, which was then colored by groups. Kruskal-Wallis rank sum tests were employed to compare taxa between groups. All statistical tests were performed in R ^24^. All plots were obtained using ggplot2 in R.

### Protein-Level Analysis of Metagenomic Data

Three publicly available gut metagenomic protein catalogs were obtained (MPA4 marker gene database, UHGP-100, and CMMG representative genomes) ^23,25,26^. The MetaPhlAn marker gene database was first converted from nucleic acid to protein sequences using Prodigal (v2.6.3) ^27^.

For each protein of interest, enzyme structures and literature were analyzed to create a rubric using methods described and employed previously (see also supplemental materials) ^28-32^.

Rubrics were employed to identify protein orthologs in all 3 databases. Resulting protein IDs were merged to their provided taxonomic annotations to create an integrated database of microbial species known to encode an enzyme of interest. For each protein, the species-level MetaPhlAn results were filtered to retain only those species that encoded the protein, and these results were summed to generate percent abundance values for each sample.

### Data Analysis

Processed data are expressed as the mean ± standard error of the mean (SEM). For the comparison between treatment groups, statistical significance was determined using a two-side t-test (Mann-Whitney test) to analyze unpaired data (**Fig. 1**). Analysis of intestinal toxicity in male and female mouse experiments (**Fig.S1**) was performed by two-way ANOVA according to sex and treatment following Tukey-Kramer’s method ^33^. For comparisons between groups, statistical significance was determined using two-side t-tests (**Fig. S1**). Comparisons of GI toxicity between groups (**Fig. 2**) were performed by two-way ANOVA following Tukey-Kramer’s method. For the comparison between treatment groups, statistical significance was determined using a two-side t-test (Mann-Whitney test) to analyze unpaired data (**Fig. 3**). Kruskal-Wallis rank sum tests were used to compare proteins of interest in metagenomic data between groups (**Fig. 4-5**). P values less than 0.05 are reported as statistically significant. All figures were generated using GraphPad Prism 10 (GraphPad Software) and R software.

**Fig. 2.**
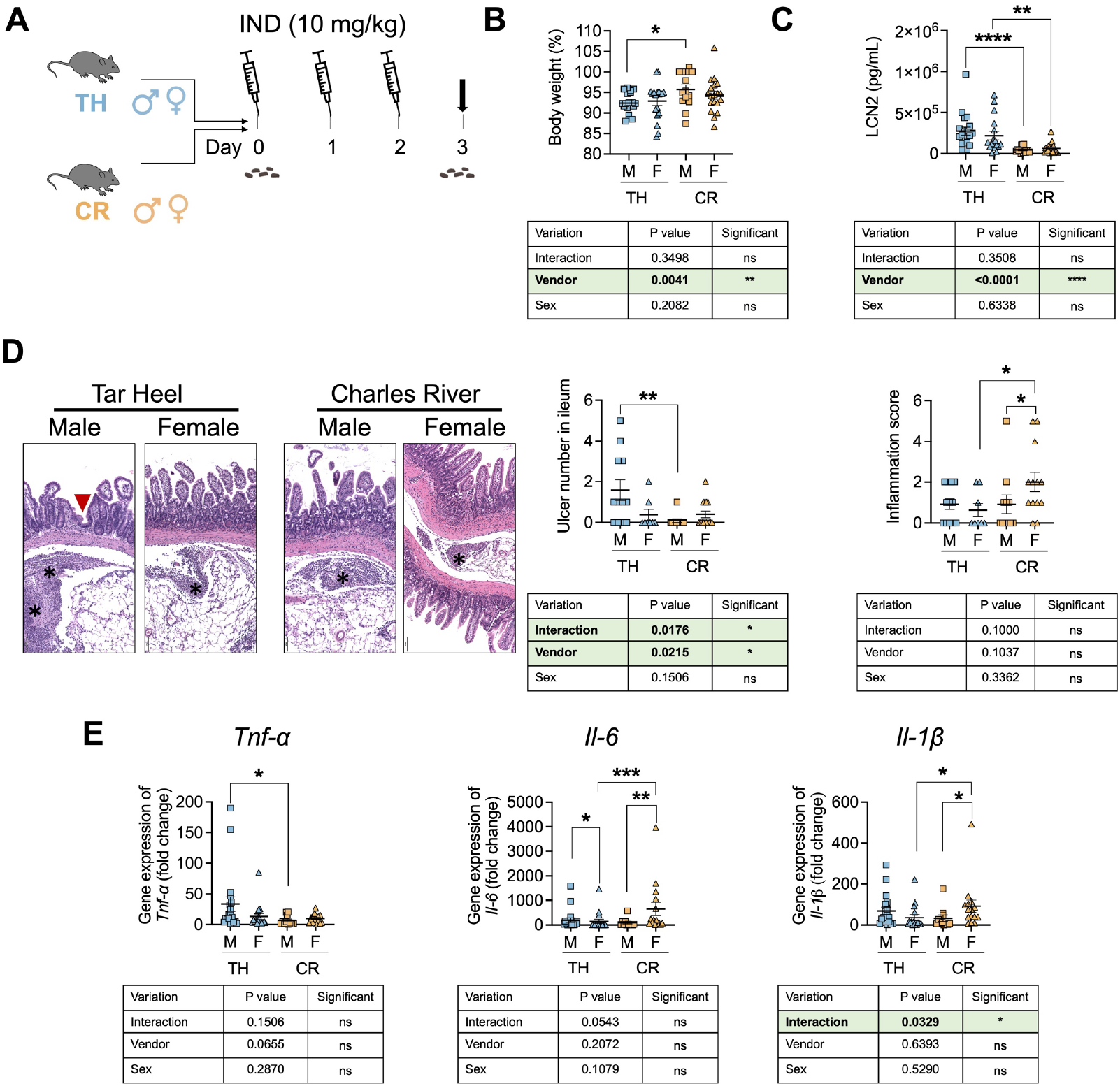
Differential indomethacin-induced GI toxicity in Tar Heel and commercial Charles River mice. **A**. Scheme of experimental design in which in-house bred Tar Heel (TH) and commercially purchased Charles River (CR) C57BL/6 male and female mice were treated with indomethacin (IND; 10 mg/kg body weight) for 3 days and then euthanized 24 h after the last dose (dark arrow). Fecal material were collected at day 0, as indicated. **B**. Percent of day 0 body weight values show male (M) CR mice lost significantly less weight than M TH mice. **C**. Fecal lipocalin-2 (LCN2) levels at day 3 are significantly higher in M and F TH mice. **D**. Differential IND-induced ulceration of the ileum as revealed by representative H&E histological images of ileum at left with an ulcer indicated with a red triangle and inflammation indicated with asterisks (scale bar = 100 μm). At right, ulcer numbers in the ileum were significantly higher in M TH mice and Significant interaction difference was found among the groups according to the vendor type and sex type by two-way ANOVA analysis. Inflammation scores in the ileum were higher in F CR mice compared to M CR and F TH mice. **E**. Gene expression of pro-inflammatory cytokines in the ileum normalized by *β-actin* show differential effects dependent on sex and vendor for *Tnf-α, Il-6*, and *Il-1β*. The GI toxicity between Tar Heel and Charles River mouse experiments was performed two-way ANOVA according to vendor type and sex type, followed by Tukey-Kramer’s method. For the comparison between two treatment groups, statistical significance was determined using two-side t-test. *P<0.05, **P<0.01, ***P<0.001, ****P<0.0001. Additional Veh and IND data for both vendors and sexes is shown in **Fig. S1**.

**Fig. 3.**
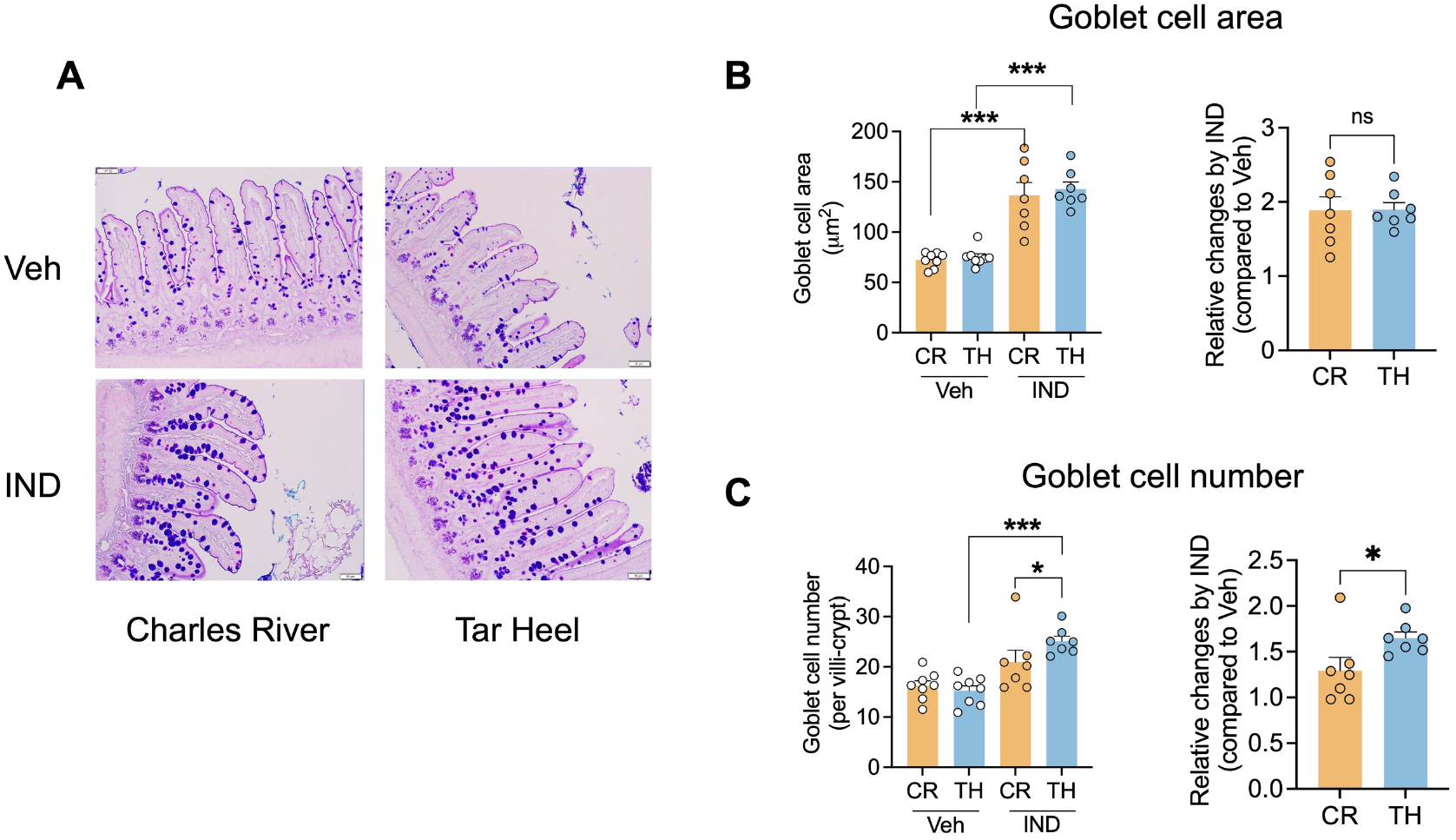
The goblet cell were increased higher in Tar Heel mice compared to Charles River mice after exposure of indomethacin. **A**. Alcian Blue/periodic acid Schiff staining of the ileal mucus in mice revealed goblet cells stained in light blue (scale bar = 50 μm). **B**. Treated with indomethacin significantly increased the goblet cell area (∼1.9 times larger compared to Veh) in mice from both vendors, with no difference observed between the vendors. **C**. Treated with indomethacin significantly increased the goblet cell number in TH mice only, with a significantly higher increase observed in TH mice compared to CR mice. CR: Charles River; TH: Tar Heel. n=7-8/group. Statistical significance was determined using two-side t-test. *P<0.05, ns: no significance.

**Fig. 4.**
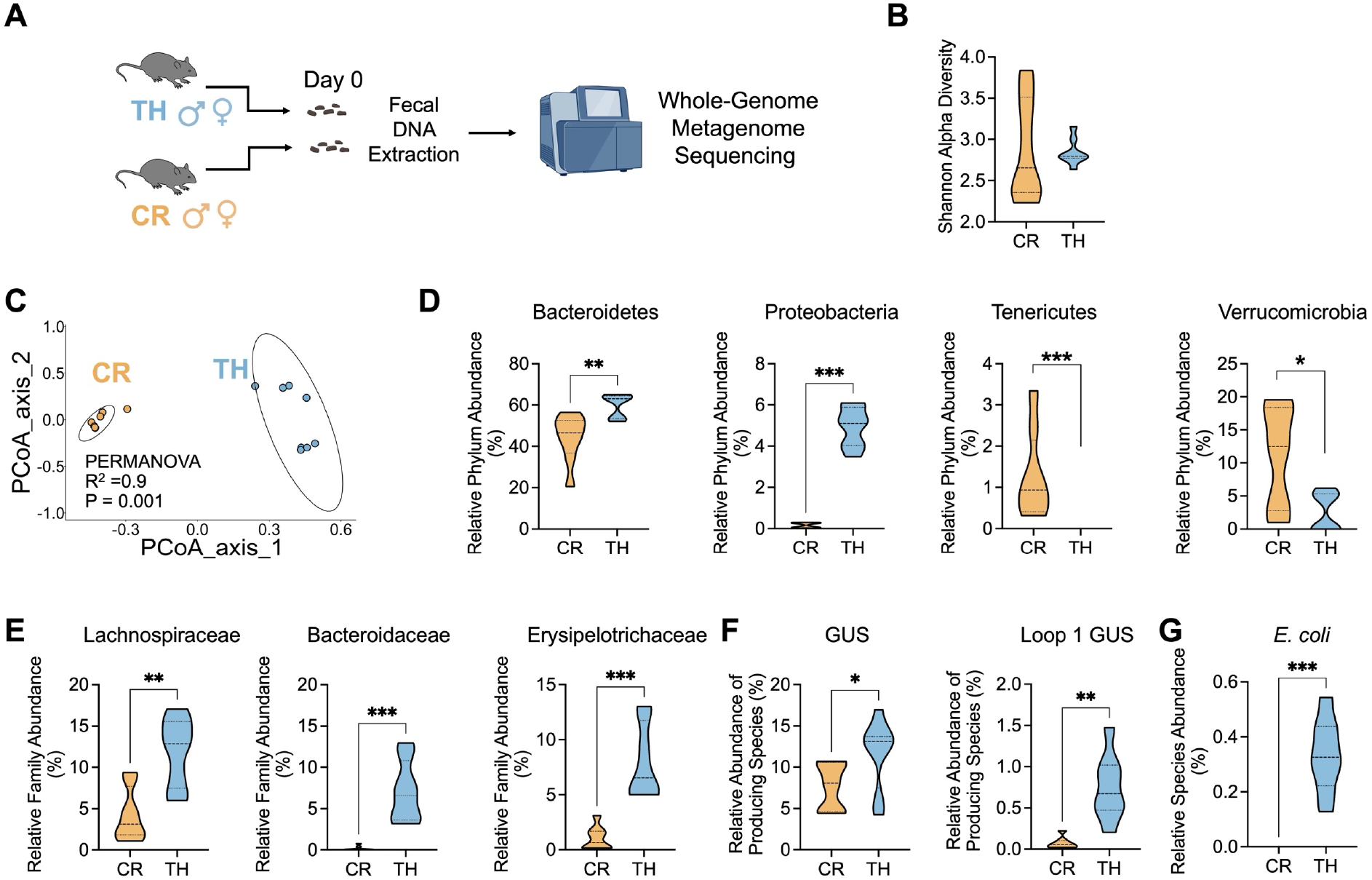
Fecal metagenomic analysis reveals altered *Bacteroidetes, Proteobacteria* abundance and β-glucuronidase (GUS) producing species in Tar Heel mice. **A**. Scheme of experimental design in which day 0 fecal samples from male and female Charles River and Tar Heel mice were examined by whole-genome metagenome sequencing. **B**. Alpha diversity did not differ between CR and TH mice. **C**. Beta diversity was significantly differed between CR mice and TH mice. **D**. At the phylum level, the abundance of Bacteroidetes and Proteobacteria were significantly higher in TH mice, while the abundance of Tenericutes and Verrucomicrobia were significantly higher in CR mice **E**. At the family level, the abundance of Lachnospiraceae, Bacteroidaceae and Erysipelotrichaceae were significantly higher in TH mice. **F-G**. GUS and Loop 1 GUS producing species were significantly higher in TH mice, as was the relative abundance of *Escherichia coli*. CR: Charles River; TH: Tar Heel. n=8/group. Kruskal-Wallis rank sum tests were used to compare all functional pathways of interest between the vendors. *P<0.05, **P<0.01, ***P<0.001, ****P<0.0001.

## RESULTS

### Indomethacin Induces Ileal Toxicity in Mice

To develop a reproducible model of NSAID-induced ileal toxicity, we treated male “Tar Heel” mice bred in-house at the University of North Carolina at Chapel Hill animal facility with indomethacin. Specifically, mice were treated with indomethacin (IND; n=8) by oral gavage at a dose of 10 mg/kg body weight or with vehicle (Veh; n=7) for three days and euthanized 24 hours after the final dose (**Fig. 1A**). Mice treated with IND exhibited significantly reduced body weight by days 2 and 3 compared to mice treated with Veh (**Fig. 1B**). IND-treated mice also exhibited significantly increased fecal lipocalin-2 levels at the end of the study compared to mice treated with Veh (**Fig. 1C**). Ileal tissues of IND-treated mice showed increased mucosal erosions and ulceration (**Fig. 1D**), and upregulated expression of the pro-inflammatory cytokines *Tnf-α, Il-6*, and *Il-1β* (**Fig. 1E**). Taken together, these results indicate that 10 mg/kg IND delivered by oral gavage for three days produces evidence of ileal toxicity in male Tar Heel mice.

### Differential Responses to Indomethacin in Tar Heel and Charles River Mice

To better understand factors that may influence NSAID-mediated intestinal injury, we next directly compared IND-treated Tar Heel and Charles River mice of both sexes (**Fig. 2A**). We found that male Tar Heel mice lost significantly more weight than male Charles River mice (**Fig. 2B**), and that all Tar Heel mice showed significantly higher fecal lipocalin-2 levels than their Charles River counterparts (**Fig. 2C**). Male Tar Heel mice exhibited significantly more ileal ulcers than male Charles River animals; conversely, female Charles River mice showed higher histologic inflammation scores than male mice from Charles River, and from female Tar Heel mice (**Fig. 2D, Fig. S1**). Finally, Tar Heel male mice also showed higher ileal *Tnf-α* expression compared to male Charles River mice, as well as higher *Il-6* expression compared to their female counterparts. Intriguingly, Charles River female mice demonstrated similar *Il-6* expression to male Tar Heel mice, which was significantly higher than their male counterparts and female Tar Heel mice. This was also the case for *Il-1β* expression (**Fig. 2E**). Taken together, these data demonstrate that mice from different breeding facilities exhibit differential toxicities and ileal damage in response to oral IND treatment. Specifically, Tar Heel mice appear to be more susceptible to IND-induced ileal ulceration than Charles River mice. A complete analysis including all vehicle groups is shown in **Fig. S1**, which further supports these findings.

Given that NSAIDs inhibit the production of prostaglandins, which are important for stimulating mucus secretion from goblet cells ^15^, we next examined the ileal goblet cells of indomethacin-treated mice using PAS-Alcian Blue staining (**Fig. 3A**). Compared to Veh groups, we found that IND treatment significantly increased ileal goblet cell area in both Charles River and Tar Heel mice (**Fig. 3B**). Moreover, IND also increased ileal goblet cell number in Tar Heel mice but not in Charles River mice (**Fig. 3C**). Overall, there were no differences in goblet cell size in Charles River and Tarheel mice treated with IND; however, IND did induce higher numbers of goblet cells in the Tarheel mice compared to their Charles River counterparts (**Fig. 3B-C**). These findings further highlight the differential response to IND in these groups.

### Tar Heel and Charles River Mice Have Distinct Intestinal Microbiomes and β-Glucuronidase Producing Taxa

Given that Tar Heel and Charles River mice display differential responses to IND, we hypothesized that their gut microbiomes may be different prior to NSAID treatment. To test this hypothesis, we performed whole-genome metagenome sequencing on fecal samples obtained at the start of the experiment, prior to vehicle or IND exposure (**Fig. 4A**). While alpha-diversity did not differ between groups (**Fig. 4B**), beta-diversity was significantly different between the Tar Heel and Charles River cohorts (**Fig. 4C**). Taxonomically, at the phylum level, we found that Tar Heel mice contained significantly higher abundances of *Bacteroidetes* and *Proteobacteria* compared to Charles River mice, while the Charles River animals contained more *Tenericutes* and *Verrucomicrobia* (**Fig. 4D, Table S2**). These differences also extended to the family level, where *Lachnospiraceae, Bacteroidaceae* and *Erysipelotrichaceae* were significantly more abundant in the Tar Heel cohort compared to Charles River mice (**Fig. 4E, Table S3**).

NSAIDs like IND are known to reach the gut as inactive glucuronides and to be reactivated by gut microbial β-glucuronidase (GUS) enzymes^34^. We have previously demonstrated the importance of β-glucuronidases in driving NSAID-induced enteropathy ^10,11^ and identified the specific molecular features of GUS responsible for carrying out this de-glucuronidation chemistry ^34^. Specifically, we have found that Loop 1 GUS are the primary class that reactivate NSAID glucuronides in vitro and in vivo. Thus, we also examined the composition of gut microbial taxa that encode GUS, including all GUS isoforms, as well as specific structural classes including Loop 1 associated with efficient drug-glucuronide processing. We found that Tar Heel mice showed significantly higher abundances of bacterial species encoding genes for all GUS enzymes and specifically for Loop 1 GUS proteins (**Fig. 4F, Table S4**). Moreover, we found that *E. coli*, a Loop 1 GUS producer, was significantly more abundant in Tar Heel compared to Charles River mice (**Fig 4G**). This is consistent with our prior findings that Loop 1 GUS are the predominant NSAID-glucuronide reactivating GUS. Taken together, these findings indicate that the Tar Heel and Charles River mice in our experiment have baseline compositional and functional differences in their gut microbiomes. Accordingly, the poor outcomes in Tar Heel mice with indomethacin (body weight, fecal lipocalin-2, and ulcers; **Fig. 2**) may have arisen due to the distinct functional capacities present in the fecal microbiomes of this cohort.

### Higher Metagenomic Abundance of Taxa Encoding Mucolytic Enzymes in Tar Heel Mice

In addition to GUS activity, we also examined other gut microbial functions that could contribute to differences between Tar Heel and Charles River mice susceptibility to IND. Given the differences noted in goblet cell numbers in these cohorts after IND-treatment, we next explored the abundances of gut microbial species that encode enzymes capable of degrading mucin polysaccharides and the mucin backbone (**Fig. 5A**). Specifically, we examined the abundances of gut microbial taxa encoding sulfatases, fucosidases, sialidases, N-acetylgalactosaminidases, N-acetylglucosaminidases, and mucinases in the fecal metagenomics data collected from our mouse cohorts. These enzyme families were chosen based on those expected to be involved in the microbial breakdown of intestinal mucus (**Fig. 5A**). We also chose four “control” enzymes expected to be present in microbes regardless of mucin metabolism: DNA polymerase (dnaPa), RNA polymerase (rpoC), enolase from glycolysis (ENO), and citrate synthase from the TCA cycle (citSyn).

**Fig. 5.**
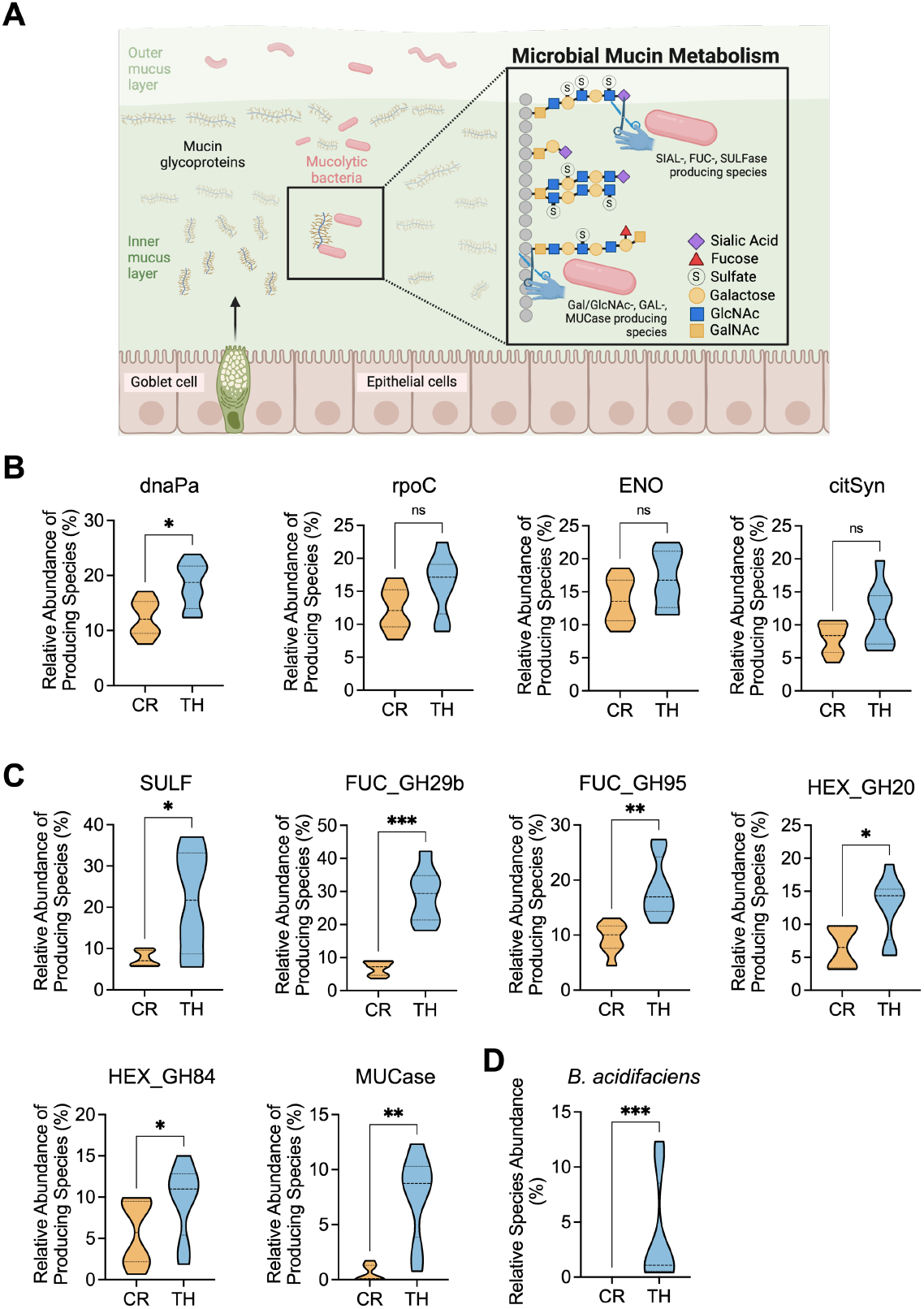
Bacterial species encoding mucolytic enzymes showed increased abundance in Tar Heel mice. **A**. Schematic of intestinal mucin polysaccharides showing sites of activity of microbial sulfatases (SULF), fucosidases (FUC), hexaminidases (HEX, including N-acetyl-glucosidases and N-aceteyl-galactosidases), and mucinases (MUCase). (Figure created with BioRender.com). **B**. The relative abundances of gut microbial species encoding “control” microbial enzymes showed no difference between the TH and CR mice for the RNA polymerase rpoC gene, the enolase (ENO) gene of glycolysis, and the citrate synthase (citSyn) of the citric acid cycle, while TH mice showed higher abundance of the DNA polymerase gene dnaPa. **C**. TH mice showed higher relative abundances of gut microbial species encoding the mucin polysaccharide degrading Sulfatase (SULF), fucosidase (FUC, both family GH29b and GH95 factors), hexosaminidase (HEX, both family GH20 N-acetyl-galactosaminidase and GH84 N-acetyl-glucosaminidase factors) and mucinase (MUCase) enzymes. **D**. The relative species abundance of the mucolytic bacteria *Bacteroides acidifaciens* was significantly higher in TH mice. CR: Charles River; TH: Tar Heel. n=8/group. Kruskal-Wallis rank sum tests were used to compare all functional pathways of interest between the vendors. *P<0.05, **P<0.01, ***P<0.001, ****P<0.0001. ns: no significance.

We found no significant differences in the abundances of microbial taxa that encode three of the four control enzymes in Tar Heel and Charles River mice (**Fig 5B**). By contrast, we found that abundances of fecal microbes encoding the following mucolytic enzymes were significantly higher in Tar Heel mice: sulfatase (SULF), fucosidases from the glycoside hydrolase (GH) families 29 and 95 (FUC_GH29, FUC_GH95), the hexosaminidase HEX) proteins N-acetylglucosaminidase (HEX_GH20) and N-acetylgalactosaminidase (HEX_GH84) and the mucinase (MUCase) enzyme (**Fig. 5C, Table S4**). Thus, the fecal microbiomes of the Tar Heel mice in our study contained a higher abundance of microbes that encoded these mucolytic factors than the Charles River mice. Finally, we found that the mucolytic species *B. acidifaciens*, which encodes each of the mucus degrading factors outlined above, was also significantly more abundant in Tar Heel than Charles River mice (**Fig. 5D**). Collectively, these results demonstrate differential abundances of specific bacterial species encoding mucolytic enzymes between Tar Heel and Charles River mice.

## DISCUSSION

In this study, we discovered significant differences in indomethacin-induced toxicity and ileal damage between mice from different facilities, either bred in-house (Tar Heel) or purchased from Charles River. Tar Heel mice exhibited enhanced ileal toxicity compared to Charles River mice, demonstrated by significantly more body weight loss, increased ulcer numbers, and elevated fecal lipocalin-2 levels. Additionally, Tar Heel mice showed a significantly higher number of goblet cells following indomethacin exposure. To investigate the mechanistic basis for these differential drug responses, we analyzed the baseline gut microbiota of both mouse populations. Whole-genome metagenomic sequencing of fecal samples revealed distinct intestinal microbiomes between Tar Heel and Charles River mice, particularly in the abundance of β-glucuronidase-producing taxa. Additional analysis also revealed higher metagenomic abundance of mucolytic enzyme-encoding taxa in Tar Heel mice. Our findings suggest that differences in intestinal microbiomes - specifically in β-glucuronidase-producing and mucolytic enzyme-producing bacteria – may contribute to the distinct intestinal toxicity responses to indomethacin between mouse populations.

The C57BL/6 mouse background is widely used to study the role of microbes in various disease models; however, differences between mouse vendors, genetic lineages and husbandry protocols have been shown to contribute to variations in phenotypes and to non-reproducibility of experimental results. A large study of the gut microbial composition by 16S rRNA sequencing of C57BL/6 mice from three different vendors (The Jackson Laboratory, Charles River Laboratories, and Taconic Biosciences) in the United States found considerable variation across eight sites, highlighting the importance of environmental conditions on microbial dynamics, which may contribute to experimental reproducibility ^18^. It was found that the microbiomes of Charles River mice were more diverse than other vendors, while the Jackson Laboratory mice changed less with time in basal mice ^18^. Our whole genome metagenomic analysis revealed fundamental differences in baseline gut microbiomes between Tar Heel and Charles River mice prior to indomethacin exposure, providing crucial insights into their differential drug responses. Distinct differences were observed in beta-diversity and specific taxonomic compositions. Tar Heel mice showed significantly higher abundances of *Bacteroidetes* and *Proteobacteria*, along with enrichments in *Lachnospiraceae, Bacteroidaceae*, and *Erysipelotrichaceae* families. An increased prevalence of *Proteobacteria* is a potential diagnostic signature of dysbiosis and risk of disease ^35^. These baseline microbiome differences suggest inherent variations in the metabolic capabilities of the gut microbiota between the two mouse populations, which could significantly influence their response to xenobiotics.

LCN-2 is an essential antimicrobial component of the innate immune system and is released mainly by mature neutrophils; this factor also serves as a critical indicator of intestinal mucosal damage and inflammation ^36,37^.A previous study showed that C57BL/6 male mice (purchased from Janvier, Le Genest St Isle, France) treated orally with indomethacin at a dose of 0.25mg/mouse (∼ 10 mg/kg body weight) for 5 days significantly increased the fecal lipocalin-2 (LCN-2) levels ^*38*^. In concordance, we also found oral indomethacin administered at 10 mg/kg body weight for 3 days significantly increased fecal LCN-2 compared to vehicle group.

Moreover, we found distinct responses between Tar Heel and Charles River mice when treating with indomethacin. Tar Heel mice demonstrated significantly greater intestinal vulnerability, characterized by elevated fecal LCN-2 levels and increased ileal ulcer numbers **(Fig. 3C**). In contrast, indomethacin exposure elicited similar increases in gene expression of proinflammatory cytokines in mice sourced from either vendor. This suggests that the differences in susceptibility to NSAID-induced enteropathy between the vendors could arise from mechanisms beyond inflammatory gene regulation, such as the microbiome or/and mucosal integrity factors.

By inhibiting cyclooxygenase enzymes, indomethacin reduces production of prostaglandins which normally protect the intestinal mucosa. Goblet cells play a crucial role in maintaining intestinal barrier function through mucus production, and their hyperplasia is often observed as a protective mechanism against intestinal injury exerted by different factors including NSAIDs^14^. This adaptive, compensatory hyperplasia response is primarily aimed at enhancing mucus production to recreate the protective barrier in the intestinal epithelium. In our study, we found that there was a differential goblet cell response to indomethacin between the vendors. While both Tar Heel and Charles River mice demonstrated increased goblet cell size following indomethacin exposure, Tar Heel mice exhibited a significantly higher number of goblet cells.

This vendor-specific difference in goblet cell hyperplasia suggests distinct host-protective responses to NSAID-induced injury. A previous study reported that treatment with indomethacin (10mg/kg) could significantly increase the goblet cell area in small intestine of C57BL/6J mice, compared to control treatment mice ^15^. The enhanced goblet cell response in Tar Heel mice may be an attempt to compensate for the increased intestinal damage they experience with indomethacin.

A key mechanism underlying NSAID-induced intestinal toxicity involves the reactivation of drug conjugates by bacterial GUS enzymes in the gut ^10,11^. Our metagenomic analysis revealed significant differences in the abundance of GUS-producing bacteria between vendors, particularly in species expressing Loop-1 GUS enzymes, which are known to be highly efficient in drug conjugate processing ^34^. Notably, the significantly higher abundance of *E. coli*, a known Loop-1 GUS producer, in Tar Heel mice may result in enhanced local reconversion of non-toxic indomethacin glucuronide conjugates to active indomethacin, contributing to increased intestinal toxicity. This enhanced capacity for drug reactivation provides a possible mechanistic explanation for the increased susceptibility to indomethacin-induced toxicity observed in Tar Heel mice. These findings align with previous studies demonstrating the critical role of bacterial GUS in NSAID enteropathy ^10,11,39^, and highlight how vendor-specific differences in microbiome composition, particularly in the abundance of bacteria capable of drug metabolism, can significantly impact experimental outcomes in pharmacological studies.

Another mechanism that may enhance NSAID-induced intestinal toxicity is an increase in microbial mucolytic enzymes, which are strongly linked to IBD pathophysiology and characterized by damage to mucosal integrity and mucus barrier dysfunction. Microbial mucolytic enzymes can break down mucus by degrading its structural components, primarily mucins ^13,40^. These enzymes play a significant role in the interaction between microbes and host mucus layers. There is evidence suggesting that IBD patients have higher levels of certain mucolytic bacteria (e.g. *Ruminococcus gnavus* and *R. torques*), contributing to mucus barrier dysfunction and exacerbation of disease symptoms ^41^. Animal models provide further insights into the mechanisms of mucolytic enzyme-induced intestinal damage. Indomethacin treatment significantly increased the abundance of the mucolytic enzyme producing taxa *Akkermansia muciniphila, Enterococcus spp*., and *Clostridium cluster* XIVa in C57BL/6 male mice (purchased from Janvier, Le Genest St Isle, France) ^*38*^. Yoshihara et al., showed that direct administration of *A. muciniphila* caused thinning of the jejunal mucus layer ^42^. In our study, we observed significantly higher abundance of the mucolytic enzyme-encoding species *B. acidifaciens* in Tar Heel mice at baseline, which may create vulnerabilities in the intestinal barrier due to increase mucus degradation. We posit a mechanism wherein enhanced microbial mucolytic activity toward intestinal mucus enhances the vulnerability of the mucosa to NSAID-induced injury, resulting in an exaggerated response to NSAID exposure. To date, no studies have compared the abundance of mucolytic enzyme-encoding species before NSAID treatment. It is possible that the relative abundance of mucus-degrading species could be a predictive biomarker for NSAID-induced intestinal damage. This may be leveraged for the clinical pain management for individuals with inflammatory bowel diseases, for whom NSAIDs are known to induce disease relapse/flares ^43^.

Beyond the distinct gut microbiome profiles, the C57BL/6 mice in this study represent two substrains—C57BL/6N (Charles River) and C57BL/6J (Tar Heel)—which may independently contribute to the differential indomethacin response. A range of phenotypic differences that have the potential to impact upon penetrance and expressivity of mutational effects between these two strains ^44^. These substrain variations may influence outcomes in metabolic models and immune responses. For example, C57BL/6J mice have a mutation of the *Nlrp12* gene; consequently, these mice exhibit a defect in neutrophil recruitment in response to a range of inflammatory stimuli compared with the C57BL/6N substrain ^45^. Further investigations are needed to understand how specific C57BL/6 substrains contribute to the NSAID response.

In conclusion, our study shows marked vendor-specific variations in NSAID-induced intestinal toxicity between mice from different breeding facilities, demonstrating the profound complexity of host-microbiome interactions in drug-related pathophysiology. Our comparative analysis between Tar Heel and Charles River mice revealed significant disparities in susceptibility to indomethacin-induced intestinal damage, characterized by marked differences in weight loss, ulcer severity, fecal LCN2 level, and ileal goblet cell morphology. Notably, despite similar inflammatory cytokine gene expression, the substantial variations in drug response suggest that the fundamental differences may lie in the bacterial microenvironment and its intricate interactions with the intestinal epithelium. Metagenomics analysis uncovered two pivotal mechanisms that may underlie these distinct drug responses: first, variations in β-glucuronidase-producing bacteria potentially modulating drug metabolism; and second, differences in mucolytic enzyme-producing bacteria that may compromise mucosal integrity. Ultimately, it is likely that several factors, including both microbiome and host differences impact the susceptibility of mice to NSAID-induced injury. These findings not only underscore the critical importance of vendor source standardization in preclinical research but also highlight the emerging paradigm of microbiome-mediated drug responses.

## ACKNOWLEDGMENTS

This research is supported by NIH/NIGMS R01s GM135218, GM137286 and GM152079 to M.R.R.; R01DK122042 to A.S.G.; R35GM155168 to A.P.B. Histological staining is supported by NIH P30 DK034987 through the Center for Gastrointestinal Biology and Disease histology core at UNC.

## AUTHOR CONTRIBUTIONS

J.Z., V.T. and M.J.Z. performed the animal experiments. J.Z and J.J.S. analyzed the data. R.S.S. quantified the pathological analysis of the intestines. J.Z., M.R.R., A.S.G. and A.P.B. wrote the manuscript. J.Z., M.R.R., A.S.G. and A.P.B. designed the study.

## DATA AVAILABILITY STATEMENT

The authors confirm that the data supporting the findings of this study are available within the article and its supplementary materials. Metagenomic sequencing data are deposited in the National Library of Medicine Sequence Read Archive with the unique identifier PRJNA1246137.

## ADDITIONAL INFORMATION

Supplementary Information is attached.

## REFERENCES

1. Wilcox, C.M., Alexander, L.N., Cotsonis, G.A. & Clark, W.S. Nonsteroidal antiinflammatory drugs are associated with both upper and lower gastrointestinal bleeding. Digestive diseases and sciences 42, 990–997 (1997).

2. Lanas, A. & Sopeña, F. Nonsteroidal anti-inflammatory drugs and lower gastrointestinal complications. Gastroenterology Clinics 38, 333–352 (2009).

3. Bindu, S., Mazumder, S. & Bandyopadhyay, U. Non-steroidal anti-inflammatory drugs (NSAIDs) and organ damage: A current perspective. Biochemical pharmacology 180, 114147 (2020).

4. Maseda, D. & Ricciotti, E. NSAID–gut microbiota interactions. Frontiers in pharmacology 11, 1153 (2020).

5. Wang, X., et al. Gut microbiota in NSAID enteropathy: new insights from inside. Frontiers in cellular and infection microbiology 11, 679396 (2021).

6. Wilson, I.D. & Nicholson, J.K. Gut microbiome interactions with drug metabolism, efficacy, and toxicity. Translational Research 179, 204–222 (2017).

7. Bhatt, A.P., et al. Nonsteroidal anti-inflammatory drug-induced leaky gut modeled using polarized monolayers of primary human intestinal epithelial cells. ACS infectious diseases 4, 46–52 (2018).

8. Ok, M.T., et al. A leaky human colon model reveals uncoupled apical/basal cytotoxicity in early Clostridioides difficile toxin exposure. American Journal of Physiology-Gastrointestinal and Liver Physiology (2023).

9. Boelsterli, U.A., Redinbo, M.R. & Saitta, K.S. Multiple NSAID-induced hits injure the small intestine: underlying mechanisms and novel strategies. toxicological sciences 131, 654–667 (2013).

10. Saitta, K.S., et al. Bacterial β-glucuronidase inhibition protects mice against enteropathy induced by indomethacin, ketoprofen or diclofenac: mode of action and pharmacokinetics. Xenobiotica 44, 28–35 (2014).

11. LoGuidice, A., Wallace, B.D., Bendel, L., Redinbo, M.R. & Boelsterli, U.A. Pharmacologic targeting of bacterial β-glucuronidase alleviates nonsteroidal anti-inflammatory drug-induced enteropathy in mice. Journal of Pharmacology and Experimental Therapeutics 341, 447–454 (2012).

12. Crittenden, S., et al. Prostaglandin E2 promotes intestinal inflammation via inhibiting microbiota-dependent regulatory T cells. Science advances 7, eabd7954 (2021).

13. Paone, P. & Cani, P.D. Mucus barrier, mucins and gut microbiota: the expected slimy partners? Gut 69, 2232–2243 (2020).

14. Gustafsson, J.K. & Johansson, M.E. The role of goblet cells and mucus in intestinal homeostasis. Nature reviews Gastroenterology & hepatology 19, 785–803 (2022).

15. Chamoun-Emanuelli, A.M., et al. NSAIDs disrupt intestinal homeostasis by suppressing macroautophagy in intestinal epithelial cells. Scientific reports 9, 14534 (2019).

16. Mukherjee, P., Roy, S., Ghosh, D. & Nandi, S. Role of animal models in biomedical research: a review. Laboratory Animal Research 38, 18 (2022).

17. McGeer, P.L. & McGeer, E.G. NSAIDs and Alzheimer disease: epidemiological, animal model and clinical studies. Neurobiology of aging 28, 639–647 (2007).

18. Long, L.L., et al. Shared and distinctive features of the gut microbiome of C57BL/6 mice from different vendors and production sites, and in response to a new vivarium. Lab animal 50, 185–195 (2021).

19. Ericsson, A.C., et al. Effects of vendor and genetic background on the composition of the fecal microbiota of inbred mice. PloS one 10, e0116704 (2015).

20. Zhang, J., et al. CYP eicosanoid pathway mediates colon cancer-promoting effects of dietary linoleic acid. The FASEB Journal 37, e23009 (2023).

21. Zhang, J., et al. Microbial enzymes induce colitis by reactivating triclosan in the mouse gastrointestinal tract. Nature communications 13, 1–14 (2022).

22. Wang, W., et al. Oxidized polyunsaturated fatty acid promotes colitis and colitis-associated tumorigenesis in mice. Journal of Crohn’s and Colitis, jjae148 (2024).

23. Blanco-Míguez, A., et al. Extending and improving metagenomic taxonomic profiling with uncharacterized species using MetaPhlAn 4. Nature Biotechnology 41, 1633–1644 (2023).

24. Team, R.C. A language and environment for statistical computing. (No Title) (2021).

25. Almeida, A., et al. A unified catalog of 204,938 reference genomes from the human gut microbiome. Nature biotechnology 39, 105–114 (2021).

26. Kieser, S., Zdobnov, E.M. & Trajkovski, M. Comprehensive mouse microbiota genome catalog reveals major difference to its human counterpart. PLOS Computational Biology 18, e1009947 (2022).

27. Hyatt, D., et al. Prodigal: prokaryotic gene recognition and translation initiation site identification. BMC bioinformatics 11, 1–11 (2010).

28. Pollet, R.M., et al. An atlas of β-glucuronidases in the human intestinal microbiome. Structure 25, 967-977. e965 (2017).

29. Simpson, J.B., et al. Metagenomics combined with activity-based proteomics point to gut bacterial enzymes that reactivate mycophenolate. Gut Microbes 14, 2107289 (2022).

30. Walker, M.E., Simpson, J.B. & Redinbo, M.R. A structural metagenomics pipeline for examining the gut microbiome. Current opinion in structural biology 75, 102416 (2022).

31. Simpson, J.B., et al. Diverse but desolate landscape of gut microbial azoreductases: A rationale for idiopathic IBD drug response. Gut Microbes 15, 2203963 (2023).

32. Simpson, J.B., et al. Gut microbial β-glucuronidases influence endobiotic homeostasis and are modulated by diverse therapeutics. Cell Host & Microbe 32, 925-944. e910 (2024).

33. Kim, H.-Y. Statistical notes for clinical researchers: Two-way analysis of variance (ANOVA)-exploring possible interaction between factors. Restorative dentistry & endodontics 39, 143–147 (2014).

34. Biernat, K.A., et al. Structure, function, and inhibition of drug reactivating human gut microbial β-glucuronidases. Scientific reports 9, 825 (2019).

35. Shin, N.-R., Whon, T.W. & Bae, J.-W. Proteobacteria: microbial signature of dysbiosis in gut microbiota. Trends in biotechnology 33, 496–503 (2015).

36. Moschen, A.R., Adolph, T.E., Gerner, R.R., Wieser, V. & Tilg, H. Lipocalin-2: a master mediator of intestinal and metabolic inflammation. Trends in Endocrinology & Metabolism 28, 388–397 (2017).

37. Moschen, A.R., et al. Lipocalin 2 protects from inflammation and tumorigenesis associated with gut microbiota alterations. Cell host & microbe 19, 455–469 (2016).

38. Yvon, S., et al. Donkey milk consumption exerts anti-inflammatory properties by normalizing antimicrobial peptides levels in Paneth’s cells in a model of ileitis in mice. European journal of nutrition 57, 155–166 (2018).

39. Yauw, S.T., et al. Microbial glucuronidase inhibition reduces severity of diclofenac-induced anastomotic leak in rats. Surgical Infections 19, 417–423 (2018).

40. McGuckin, M.A., Lindén, S.K., Sutton, P. & Florin, T.H. Mucin dynamics and enteric pathogens. Nature Reviews Microbiology 9, 265–278 (2011).

41. Png, C.W., et al. Mucolytic bacteria with increased prevalence in IBD mucosa augmentin vitroutilization of mucin by other bacteria. Official journal of the American College of Gastroenterology| ACG 105, 2420–2428 (2010).

42. Yoshihara, T., et al. The protective effect of Bifidobacterium bifidum G9-1 against mucus degradation by Akkermansia muciniphila following small intestine injury caused by a proton pump inhibitor and aspirin. Gut Microbes 11, 1385–1404 (2020).

43. Long, M.D., et al. Role of nonsteroidal anti-inflammatory drugs in exacerbations of inflammatory bowel disease. Journal of clinical gastroenterology 50, 152–156 (2016).

44. Simon, M.M., et al. A comparative phenotypic and genomic analysis of C57BL/6J and C57BL/6N mouse strains. Genome biology 14, 1–22 (2013).

45. Ulland, T.K., et al. Nlrp12 mutation causes C57BL/6J strain-specific defect in neutrophil recruitment. Nature communications 7, 13180 (2016).

