## Supplementary Information for "Commercially Purchased and In-House Bred C57BL/6 Mice with Different Gut Microbiota Exhibit Distinct Indomethacin-Induced Toxicities"

### SUPPORTING INFORMATION

**Table S1. Sequences of primers in qRT-PCR.**

| Gene | Forward primer | Reverse primer |
|---|---|---|
| *β-actin* | AGCCATGTACGTAGCCATCCAG | TGGCGTGAGGGAGAGCATAG |
| *Tnf-α* | ACCCTCACACTCAGATCATCTTCTC | TGAGATCCATGCCGTTGG |
| *Il-6* | TACCACTTCACAAGTCGGAGGC | CTGCAAGTGCATCATCGTTGTTC |
| *Il-1β* | GTGGACCTTCCAGGATGAGG | CGGAGCCTGTAGTGCAGTTG |

**Table S2. Comparison of the phylum-level differences in abundance of fecal microbiota between Charles River and Tar Heel mice**. Here, Z< 0 means the abundance is higher in Tar Heel vs. Charles River mice. P<0.05. n=8/group. The names of the phyla with significant differences and their associated P values are shown in bold.

| Phylum | Min | Q1 | Median | Q3 | Max | Mean | SD | IQR | Z | P value |
|---|---|---|---|---|---|---|---|---|---|---|
| ***Bacteroidetes*** | 20.6132 | 46.5615 | 53.0505 | 63.0742 | 64.9889 | 52.0062 | 12.0661 | 15.0362 | -2.9406 | **0.0033** |
| *Firmicutes* | 19.4788 | 24.9587 | 26.5945 | 33.8707 | 52.2457 | 30.3113 | 8.9437 | 7.9631 | 0.9452 | 0.3446 |
| ***Verrucomicrobia*** | 0.0000 | 0.4925 | 5.0818 | 12.4917 | 19.5549 | 6.9217 | 7.0466 | 11.5221 | 2.3276 | **0.0199** |
| *Unclassified* | 0.5806 | 2.7841 | 4.9965 | 5.9464 | 21.8982 | 5.9067 | 5.2378 | 2.9025 | 0.5251 | 0.5995 |
| ***Proteobacteria*** | 0.0263 | 0.1669 | 1.8918 | 5.1081 | 6.0898 | 2.5462 | 2.5464 | 4.8507 | -3.3607 | **0.0008** |
| *Actinobacteria* | 0.1809 | 0.5181 | 0.8688 | 1.3942 | 2.3383 | 0.9785 | 0.5971 | 0.7386 | -1.0502 | 0.2936 |
| *Deferribacteres* | 0.0217 | 0.1739 | 0.4778 | 1.0221 | 2.8037 | 0.6850 | 0.6861 | 0.8394 | -0.4201 | 0.6744 |
| ***Tenericutes*** | 0.0000 | 0.0000 | 0.1571 | 0.9354 | 3.3419 | 0.6389 | 0.9865 | 0.8769 | 3.5897 | **0.0003** |
| *Euryarchaeota* | 0.0000 | 0.0000 | 0.0000 | 0.0000 | 0.0550 | 0.0056 | 0.0158 | 0.0000 | -1.4606 | 0.1441 |

**Table S3. Comparison of the family-level differences in abundance of fecal microbiota between Charles River and Tar Heel mice**. Here, rows with Z< 0 are colored orange and indicate that the abundance is higher in Tar Heel vs. Charles River mice, while rows with Z> 0 are colored green and indicate that the abundance is higher in Charles River vs. Tar Heel mice,. P<0.05. n=8/group.

| Taxonomy to Family | Min | Q1 | Median | Q3 | Max | Mean | SD | IQR | Z | P value |
|---|---|---|---|---|---|---|---|---|---|---|
| *Proteobacteria__Betaproteobacteria__Burkholderiales__Sutterellaceae* | 0.0000 | 0.0000 | 0.6521 | 2.4070 | 2.7511 | 1.0768 | 1.1766 | 2.3901 | -3.5897 | 0.0003 |
| *Firmicutes__Bacilli__Lactobacillales__Streptococcaceae* | 0.0000 | 0.0000 | 0.1657 | 0.9162 | 1.4939 | 0.4595 | 0.5530 | 0.8831 | -3.5897 | 0.0003 |
| *Bacteroidetes__Bacteroidia__Bacteroidales__Barnesiellaceae* | 0.0000 | 0.0000 | 0.1495 | 5.1941 | 14.2369 | 2.9889 | 5.1305 | 3.0289 | -3.5897 | 0.0003 |
| *Actinobacteria__Coriobacteriia__Coriobacteriales__Atopobiaceae* | 0.0000 | 0.0000 | 0.0905 | 1.1312 | 1.7599 | 0.4828 | 0.6623 | 1.1237 | -3.5897 | 0.0003 |
| *Proteobacteria__Epsilonproteobacteria__Campylobacterales__Helicobacteraceae* | 0.0000 | 0.0000 | 0.0804 | 0.8828 | 2.5910 | 0.5752 | 0.9071 | 0.6493 | -3.5897 | 0.0003 |
| *Firmicutes__Bacilli__Lactobacillales__Enterococcaceae* | 0.0000 | 0.0000 | 0.0762 | 0.3022 | 0.5889 | 0.1696 | 0.1969 | 0.2983 | -3.5897 | 0.0003 |
| *Firmicutes__CFGB2833__OFGB2833__FGB2833* | 0.0000 | 0.0000 | 0.0718 | 0.5529 | 1.6125 | 0.3507 | 0.5173 | 0.4693 | -3.5897 | 0.0003 |
| *Deferribacteres__CFGB75907__OFGB75907__FGB75907* | 0.0000 | 0.0000 | 0.0109 | 0.8016 | 2.7942 | 0.4263 | 0.7521 | 0.6965 | -3.5897 | 0.0003 |
| *Firmicutes__CFGB9637__OFGB9637__FGB9637* | 0.0000 | 0.0000 | 0.0039 | 0.0279 | 0.1450 | 0.0223 | 0.0375 | 0.0261 | -3.5897 | 0.0003 |
| *Proteobacteria__Gammaproteobacteria__Enterobacterales__Enterobacteriaceae* | 0.0000 | 0.0000 | 0.0641 | 0.3262 | 0.5435 | 0.1665 | 0.1948 | 0.3172 | -3.5082 | 0.0005 |
| *Bacteroidetes__Bacteroidia__Bacteroidales__Bacteroidaceae* | 0.0000 | 0.0010 | 1.9611 | 6.5606 | 12.9222 | 3.6119 | 4.3828 | 6.2413 | -3.3857 | 0.0007 |
| *Firmicutes__Erysipelotrichia__Erysipelotrichales__Erysipelotrichaceae* | 0.1562 | 0.6535 | 4.0547 | 6.5374 | 13.0062 | 4.4677 | 4.2868 | 5.2524 | -3.3607 | 0.0008 |
| *Firmicutes__Clostridia__Eubacteriales__Lachnospiraceae* | 1.1117 | 3.1190 | 7.7879 | 12.8372 | 17.0564 | 8.0846 | 5.3453 | 9.0540 | -2.9406 | 0.0033 |
| *Bacteroidetes__Bacteroidia__Bacteroidales__Rikenellaceae* | 0.0000 | 0.0000 | 0.0000 | 0.0037 | 0.0441 | 0.0071 | 0.0150 | 0.0030 | -2.5538 | 0.0107 |
| *Bacteroidetes__CFGB4425__OFGB4425__FGB4425* | 0.0000 | 0.0000 | 0.0000 | 0.1996 | 1.7179 | 0.2626 | 0.5513 | 0.0998 | -2.2075 | 0.0273 |

| | | | | | | | | | | |
|---|---|---|---|---|---|---|---|---|---|---|
| *Firmicutes__CFGB10239__OFGB10239__FGB10239* | 0.0000 | 0.0000 | 0.0000 | 0.0362 | 0.2593 | 0.0445 | 0.0896 | 0.0181 | -2.2075 | 0.0273 |
| *Bacteroidetes__CFGB76719__OFGB76719__FGB76719* | 0.0000 | 0.0000 | 0.0000 | 6.5476 | 20.3938 | 3.9814 | 7.2600 | 3.2738 | -2.2075 | 0.0273 |
| *Actinobacteria__CFGB4875__OFGB4875__FGB4875* | 0.0000 | 0.0000 | 0.0000 | 0.0244 | 1.4565 | 0.1151 | 0.3637 | 0.0122 | -2.2075 | 0.0273 |
| *Bacteroidetes__CFGB76294__OFGB76294__FGB76294* | 0.0000 | 0.0000 | 0.0000 | 0.0158 | 1.0499 | 0.0785 | 0.2609 | 0.0079 | -2.2075 | 0.0273 |
| *Proteobacteria__CFGB76450__OFGB76450__FGB76450* | 0.0000 | 0.0000 | 0.0000 | 0.1848 | 0.5991 | 0.1187 | 0.2167 | 0.0924 | -2.2075 | 0.0273 |
| *Proteobacteria__CFGB76404__OFGB76404__FGB76404* | 0.0000 | 0.0000 | 0.0000 | 0.2652 | 2.4311 | 0.3147 | 0.6784 | 0.1326 | -2.2075 | 0.0273 |
| *Firmicutes__CFGB9495__OFGB9495__FGB9495* | 0.0000 | 0.0000 | 0.0000 | 0.1970 | 4.4214 | 0.5380 | 1.2581 | 0.0985 | -2.2075 | 0.0273 |
| *Bacteroidetes__CFGB76349__OFGB76349__FGB76349* | 0.0000 | 0.0000 | 0.0000 | 0.1011 | 0.4835 | 0.0863 | 0.1640 | 0.0506 | -2.2075 | 0.0273 |
| *Firmicutes__CFGB76576__OFGB76576__FGB76576* | 0.0000 | 0.0000 | 0.0000 | 0.6434 | 2.0038 | 0.3907 | 0.7124 | 0.3217 | 2.2075 | 0.0273 |
| *Firmicutes__CFGB76355__OFGB76355__FGB76355* | 0.0000 | 0.0000 | 0.0000 | 0.6048 | 1.7107 | 0.3546 | 0.6413 | 0.3024 | 2.2075 | 0.0273 |
| *Firmicutes__CFGB9589__OFGB9589__FGB9589* | 0.0000 | 0.0000 | 0.0000 | 0.1078 | 0.4698 | 0.0843 | 0.1610 | 0.0539 | 2.2075 | 0.0273 |
| *Firmicutes__CFGB9827__OFGB9827__FGB9827* | 0.0000 | 0.0000 | 0.0000 | 0.0373 | 0.0939 | 0.0219 | 0.0393 | 0.0187 | 2.2075 | 0.0273 |
| *Firmicutes__Clostridia__Eubacteriales__Eubacteriales_Family_XIII_Incertae_Sedis* | 0.0000 | 0.0000 | 0.0000 | 0.0247 | 0.1136 | 0.0195 | 0.0371 | 0.0123 | 2.2075 | 0.0273 |
| *Firmicutes__CFGB1227__OFGB1227__FGB1227* | 0.0000 | 0.0000 | 0.0000 | 0.0055 | 0.0329 | 0.0063 | 0.0121 | 0.0027 | 2.2075 | 0.0273 |
| *Firmicutes__CFGB9479__OFGB9479__FGB9479* | 0.0000 | 0.0000 | 0.0000 | 0.0089 | 0.0301 | 0.0056 | 0.0104 | 0.0045 | 2.2075 | 0.0273 |
| *Firmicutes__CFGB75986__OFGB75986__FGB75986* | 0.0000 | 0.0000 | 0.0000 | 0.0001 | 0.0021 | 0.0002 | 0.0006 | 0.0000 | 2.2075 | 0.0273 |
| *Firmicutes__CFGB76586__OFGB76586__FGB76586* | 0.0000 | 0.0000 | 0.0000 | 0.6036 | 1.7968 | 0.3781 | 0.6876 | 0.3018 | 2.2075 | 0.0273 |
| *Firmicutes__CFGB76536__OFGB76536__FGB76536* | 0.0000 | 0.0000 | 0.0000 | 0.1828 | 2.3772 | 0.2762 | 0.6532 | 0.0914 | 2.2075 | 0.0273 |
| *Actinobacteria__Actinomycetia__Micrococcales__Brevibacteriaceae* | 0.0000 | 0.0000 | 0.0000 | 0.0173 | 0.5351 | 0.0855 | 0.1825 | 0.0087 | 2.2075 | 0.0273 |

| | | | | | | | | | | |
|---|---|---|---|---|---|---|---|---|---|---|
| *Firmicutes__CFGB76166__OFGB76166__FGB76166* | 0.0000 | 0.0000 | 0.0000 | 0.0500 | 0.3048 | 0.0540 | 0.1065 | 0.0250 | 2.2075 | 0.0273 |
| *Firmicutes__Clostridia__Eubacteriales__Eubacteriales_unclassified* | 0.0000 | 0.0000 | 0.0000 | 0.0917 | 0.3106 | 0.0566 | 0.1046 | 0.0458 | 2.2075 | 0.0273 |
| *Firmicutes__CFGB1292__OFGB1292__FGB1292* | 0.0000 | 0.0000 | 0.0000 | 0.0047 | 0.0355 | 0.0058 | 0.0114 | 0.0023 | 2.2075 | 0.0273 |
| *Firmicutes__CFGB3297__OFGB3297__FGB3297* | 0.0000 | 0.0000 | 0.0000 | 0.0024 | 0.0085 | 0.0015 | 0.0028 | 0.0012 | 2.2075 | 0.0273 |
| *Bacteroidetes__CFGB74567__OFGB74567__FGB74567* | 0.0000 | 0.0000 | 0.0000 | 0.0000 | 0.0004 | 0.0001 | 0.0001 | 0.0000 | 2.2075 | 0.0273 |
| *Verrucomicrobia__Verrucomicrobiae__Verrucomicrobiales__Akkermansiaceae* | 0.0000 | 0.4925 | 5.0818 | 12.4917 | 19.5549 | 6.9217 | 7.0466 | 11.5221 | 2.3276 | 0.0199 |
| *Bacteroidetes__Bacteroidia__Bacteroidales__Tannerellaceae* | 0.1751 | 0.8828 | 1.8513 | 6.3223 | 7.3177 | 3.0339 | 2.7873 | 4.9863 | 2.4155 | 0.0157 |
| *Firmicutes__Clostridia__Eubacteriales__Oscillospiraceae* | 0.0110 | 0.3493 | 1.0848 | 3.0347 | 11.2897 | 2.3613 | 3.1693 | 2.3305 | 2.6255 | 0.0087 |
| *Firmicutes__CFGB37314__OFGB37314__FGB37314* | 0.0000 | 0.0256 | 0.3278 | 0.8134 | 2.4196 | 0.6297 | 0.8012 | 0.7400 | 2.6450 | 0.0082 |
| *Firmicutes__CFGB1494__OFGB1494__FGB1494* | 0.0000 | 0.0000 | 0.0000 | 0.0039 | 0.6449 | 0.0861 | 0.2062 | 0.0028 | 2.8963 | 0.0038 |
| *Firmicutes__CFGB76071__OFGB76071__FGB76071* | 0.0000 | 0.0000 | 0.0000 | 0.0200 | 0.0581 | 0.0124 | 0.0217 | 0.0115 | 2.8963 | 0.0038 |
| *Firmicutes__Erysipelotrichia__Erysipelotrichales__Turicibacteraceae* | 0.0000 | 0.0000 | 0.0000 | 0.3476 | 0.6502 | 0.1568 | 0.2226 | 0.3383 | 3.2404 | 0.0012 |
| *Firmicutes__CFGB48875__OFGB48875__FGB48875* | 0.0000 | 0.0000 | 0.0000 | 0.0950 | 0.1658 | 0.0439 | 0.0562 | 0.0937 | 3.2404 | 0.0012 |
| *Bacteria_unclassified__CFGB2838__OFGB2838__FGB2838* | 0.0000 | 0.0000 | 0.0000 | 0.0492 | 0.1733 | 0.0333 | 0.0555 | 0.0405 | 3.2404 | 0.0012 |
| *Firmicutes__Bacilli__Lactobacillales__Lactobacillaceae* | 0.3309 | 1.2276 | 3.4793 | 5.7916 | 8.8879 | 3.6690 | 2.7024 | 4.2029 | 3.3607 | 0.0008 |
| *Firmicutes__Tissierellia__Tissierellales__Peptoniphilaceae* | 0.0051 | 0.0096 | 0.2865 | 1.7329 | 3.5356 | 0.9784 | 1.2461 | 1.5191 | 3.3607 | 0.0008 |
| *Deferribacteres__Deferribacteres__Deferribacterales__Deferribacteraceae* | 0.0000 | 0.0001 | 0.0760 | 0.3621 | 1.1473 | 0.2587 | 0.3693 | 0.3609 | 3.3857 | 0.0007 |
| *Bacteria_unclassified__CFGB76064__OFGB76064__FGB76064* | 0.0000 | 0.0000 | 0.2854 | 0.9515 | 6.8521 | 1.2086 | 2.0631 | 0.9317 | 3.4506 | 0.0006 |
| *Bacteria_unclassified__CFGB76313__OFGB76313__FGB76313* | 0.0000 | 0.0000 | 0.0652 | 0.5756 | 3.5120 | 0.5915 | 1.0708 | 0.3899 | 3.5082 | 0.0005 |

| | | | | | | | | | | |
|---|---|---|---|---|---|---|---|---|---|---|
| *Bacteroidetes__CFGB1791__OFGB1791__FGB1791* | 0.0000 | 0.0000 | 0.1571 | 0.9354 | 3.3419 | 0.6389 | 0.9865 | 0.8769 | 3.5897 | 0.0003 |
| *Firmicutes__Clostridia__Eubacteriales__Christensenellaceae* | 0.0000 | 0.0000 | 0.1546 | 0.7180 | 1.9991 | 0.5001 | 0.6927 | 0.6702 | 3.5897 | 0.0003 |
| *Firmicutes__Clostridia__Clostridia_unclassified__Clostridia_unclassified* | 0.0000 | 0.0000 | 0.1241 | 0.9412 | 1.4808 | 0.5021 | 0.5876 | 0.9091 | 3.5897 | 0.0003 |
| *Firmicutes__CFGB6645__OFGB6645__FGB6645* | 0.0000 | 0.0000 | 0.1054 | 1.0277 | 1.8173 | 0.4728 | 0.6589 | 0.8466 | 3.5897 | 0.0003 |
| *Firmicutes__CFGB76074__OFGB76074__FGB76074* | 0.0000 | 0.0000 | 0.0984 | 0.4864 | 1.0344 | 0.2844 | 0.3625 | 0.4389 | 3.5897 | 0.0003 |
| *Candidatus_Saccharibacteria__CFGB8157__OFGB8157__FGB8157* | 0.0000 | 0.0000 | 0.0976 | 1.3896 | 2.4561 | 0.6332 | 0.8143 | 1.3893 | 3.5897 | 0.0003 |
| *Firmicutes__CFGB10344__OFGB10344__FGB10344* | 0.0000 | 0.0000 | 0.0707 | 0.8874 | 2.0249 | 0.4891 | 0.6414 | 0.8784 | 3.5897 | 0.0003 |
| *Firmicutes__Clostridia__Eubacteriales__Clostridiaceae* | 0.0000 | 0.0000 | 0.0655 | 0.6473 | 1.1946 | 0.3227 | 0.4215 | 0.6028 | 3.5897 | 0.0003 |
| *Firmicutes__CFGB76293__OFGB76293__FGB76293* | 0.0000 | 0.0000 | 0.0408 | 0.3103 | 0.8893 | 0.1826 | 0.2826 | 0.2143 | 3.5897 | 0.0003 |
| *Actinobacteria__Acidimicrobiia__Acidimicrobiales__Ilumatobacteraceae* | 0.0000 | 0.0000 | 0.0380 | 0.1613 | 0.4978 | 0.1122 | 0.1558 | 0.1406 | 3.5897 | 0.0003 |
| *Firmicutes__CFGB49426__OFGB49426__FGB49426* | 0.0000 | 0.0000 | 0.0331 | 0.1607 | 0.3118 | 0.0837 | 0.1047 | 0.1477 | 3.5897 | 0.0003 |
| *Firmicutes__CFGB8152__OFGB8152__FGB8152* | 0.0000 | 0.0000 | 0.0302 | 0.2330 | 2.0655 | 0.2364 | 0.5214 | 0.1633 | 3.5897 | 0.0003 |
| *Firmicutes__Clostridia__Clostridiales__Peptostreptococcaceae* | 0.0000 | 0.0000 | 0.0290 | 0.5742 | 1.1029 | 0.2888 | 0.4479 | 0.3843 | 3.5897 | 0.0003 |
| *Firmicutes__CFGB28980__OFGB28980__FGB28980* | 0.0000 | 0.0000 | 0.0169 | 0.1041 | 1.2323 | 0.1470 | 0.3239 | 0.0802 | 3.5897 | 0.0003 |
| *Firmicutes__CFGB28820__OFGB28820__FGB28820* | 0.0000 | 0.0000 | 0.0160 | 0.1978 | 0.5313 | 0.1086 | 0.1586 | 0.1847 | 3.5897 | 0.0003 |
| *Firmicutes__CFGB76144__OFGB76144__FGB76144* | 0.0000 | 0.0000 | 0.0074 | 0.0467 | 0.1233 | 0.0270 | 0.0378 | 0.0443 | 3.5897 | 0.0003 |
| *Firmicutes__CFGB76351__OFGB76351__FGB76351* | 0.0000 | 0.0000 | 0.0049 | 0.2135 | 0.5823 | 0.1244 | 0.1939 | 0.1808 | 3.5897 | 0.0003 |
| *Firmicutes__CFGB76790__OFGB76790__FGB76790* | 0.0000 | 0.0000 | 0.0021 | 0.0789 | 0.3364 | 0.0687 | 0.1236 | 0.0458 | 3.5897 | 0.0003 |
| *Firmicutes__CFGB76234__OFGB76234__FGB76234* | 0.0000 | 0.0000 | 0.0020 | 0.0162 | 0.0511 | 0.0096 | 0.0145 | 0.0153 | 3.5897 | 0.0003 |

| | | | | | | | | | | |
|---|---|---|---|---|---|---|---|---|---|---|
| *Firmicutes__CFGB76077__OFGB76077__FGB76077* | 0.0000 | 0.0000 | 0.0019 | 0.0513 | 0.3011 | 0.0541 | 0.0988 | 0.0357 | 3.5897 | 0.0003 |
| *Firmicutes__CFGB76308__OFGB76308__FGB76308* | 0.0000 | 0.0000 | 0.0011 | 0.0069 | 0.0560 | 0.0093 | 0.0171 | 0.0060 | 3.5897 | 0.0003 |
| *Firmicutes__CFGB9633__OFGB9633__FGB9633* | 0.0000 | 0.0000 | 0.0009 | 0.0221 | 1.4290 | 0.2109 | 0.4557 | 0.0159 | 3.5897 | 0.0003 |

**Table S4. Relative abundance differences in gut microbial taxa capable of expressing the indicated genes between Tar Heel and Charles River mice**. Here, Z< 0 means the abundance is higher in Tar Heel vs. Charles River mice. P<0.05. n=8/group. The names of the genes with significant differences and their associated P values are shown in bold

| Gene | Min | Q1 | Median | Q3 | Max | Mean | SD | IQR | Z | P value |
|---|---|---|---|---|---|---|---|---|---|---|
| **dnaPa** | 7.5299 | 11.4467 | 15.0760 | 18.7980 | 23.8295 | 15.3094 | 4.8411 | 6.0389 | -2.3105 | **0.0209** |
| rpoC | 7.6724 | 10.4346 | 14.7470 | 17.5996 | 22.3836 | 14.2045 | 4.2875 | 6.8575 | -1.6803 | 0.0929 |
| ENO | 8.9898 | 11.6472 | 15.1755 | 18.4353 | 22.4497 | 15.3463 | 4.1168 | 6.6701 | -1.5753 | 0.1152 |
| citSyn | 4.3104 | 6.5735 | 8.6310 | 11.7968 | 19.6922 | 9.6383 | 4.0328 | 4.4284 | -1.4703 | 0.1415 |
| **GUS** | 4.2435 | 5.3486 | 10.4836 | 13.1273 | 16.9214 | 9.6629 | 4.0437 | 7.3698 | -1.9954 | **0.0460** |
| **L1_GUS** | 0.0174 | 0.0537 | 0.2131 | 0.6720 | 1.4733 | 0.4109 | 0.4463 | 0.5560 | -3.2557 | **0.0011** |
| **CTD_GUS** | 0.0000 | 0.2569 | 1.0235 | 1.7657 | 2.6007 | 1.0628 | 0.8750 | 1.4282 | 2.9428 | **0.0033** |
| **L1_CTD_GUS** | 0.2036 | 0.5492 | 1.1942 | 1.8164 | 2.7112 | 1.2653 | 0.7801 | 1.2541 | 2.4155 | **0.0157** |
| L2_GUS | 0.4526 | 1.2747 | 2.3762 | 5.1820 | 7.2025 | 3.1799 | 2.5104 | 2.8458 | 1.7854 | 0.0742 |
| mL1_GUS | 0.6100 | 2.4158 | 3.9461 | 7.0902 | 9.9643 | 4.6231 | 2.7965 | 4.5346 | 0.0000 | 1.0000 |
| NL_GUS | 1.0425 | 3.6155 | 8.5286 | 10.5406 | 12.7551 | 7.3061 | 3.7188 | 6.5360 | -1.8904 | 0.0587 |
| SIAL | 3.7197 | 7.5363 | 10.9617 | 13.4250 | 19.5005 | 10.7908 | 4.2052 | 5.0455 | -1.5753 | 0.1152 |
| **SULF** | 5.5429 | 6.2392 | 9.4721 | 21.7029 | 37.0226 | 14.5595 | 11.1288 | 12.4597 | -2.1004 | **0.0357** |
| FUC_GH29a | 0.0000 | 0.3050 | 4.1063 | 7.6058 | 8.8516 | 4.1350 | 3.4981 | 6.9565 | 1.5870 | 0.1125 |
| **FUC_GH29b** | 3.6367 | 7.2252 | 13.5766 | 29.4332 | 42.1483 | 17.7814 | 12.7549 | 20.9604 | -3.3607 | **0.0008** |
| **FUC_GH95** | 4.4558 | 10.0744 | 12.6204 | 16.9726 | 27.3575 | 14.0058 | 6.2440 | 6.0332 | -3.2557 | **0.0011** |
| bGAL_GH2 | 6.2477 | 8.8019 | 13.4837 | 16.3208 | 21.5323 | 13.1043 | 4.5018 | 7.3120 | -1.1552 | 0.2480 |
| bGAL_GH35 | 6.2991 | 8.5697 | 11.4914 | 17.6060 | 22.9899 | 12.9252 | 5.2457 | 8.7854 | -1.8904 | 0.0587 |
| bGAL_GH42 | 0.2273 | 0.9237 | 1.2772 | 2.1784 | 5.3397 | 1.6361 | 1.2405 | 1.1649 | -1.5753 | 0.1152 |
| **hexAm_GH20** | 3.1819 | 5.2559 | 9.5440 | 14.3160 | 19.0565 | 9.6240 | 5.0073 | 9.0019 | -2.5205 | **0.0117** |
| **hexAm_GH84** | 0.6680 | 2.6962 | 9.3261 | 10.9954 | 14.9876 | 7.6639 | 4.5765 | 7.8294 | -1.9954 | **0.0460** |
| **MUCase** | 0.0000 | 0.1046 | 1.5394 | 8.7463 | 12.3075 | 4.0792 | 4.5558 | 8.5364 | -3.0683 | **0.0022** |

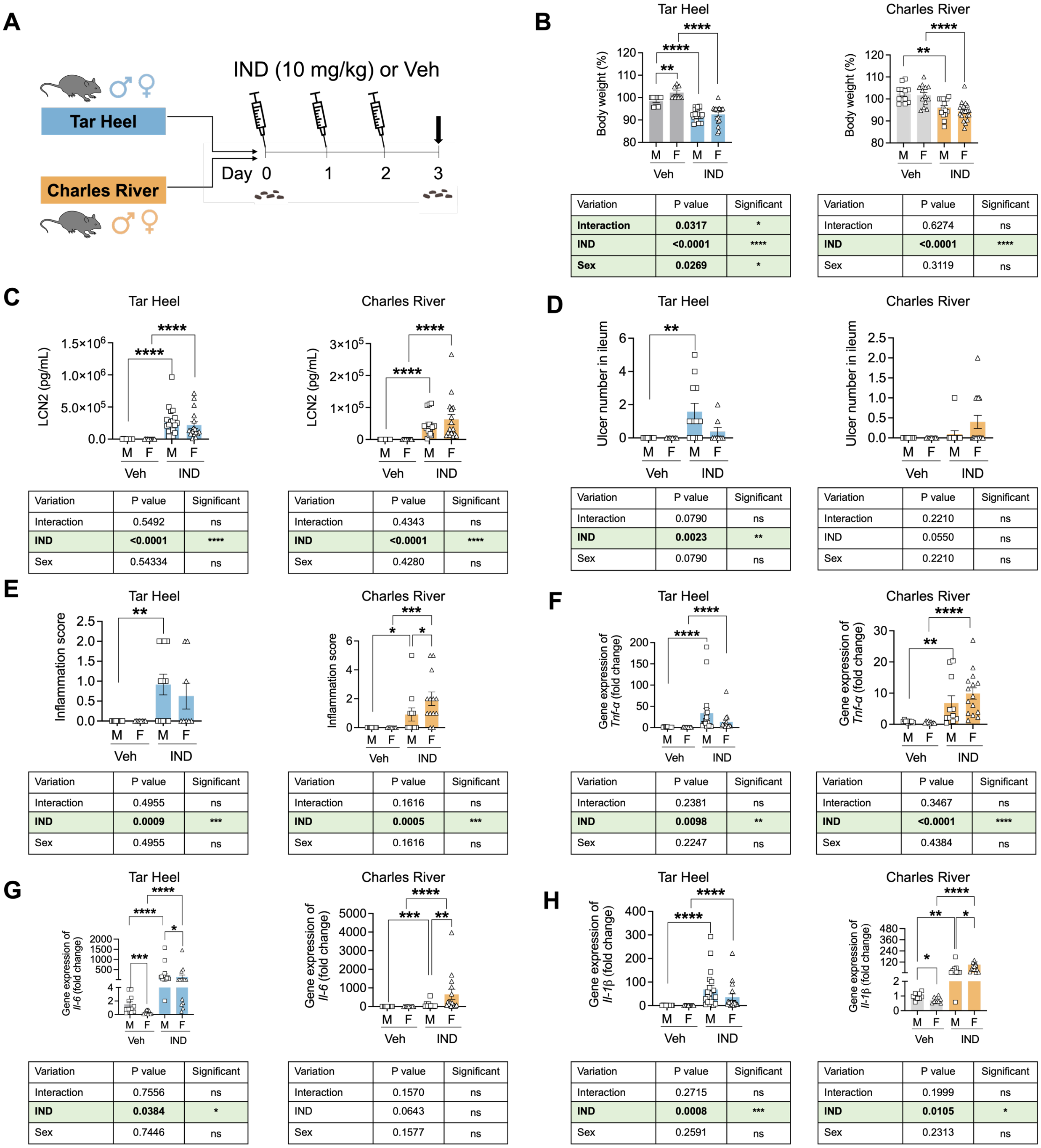


**B. Tar Heel**

| Variation | P value | Significant |
|---|---|---|
| **Interaction** | **0.0317** | * |
| **IND** | **<0.0001** | **** |
| **Sex** | **0.0269** | * |

**B. Charles River**

| Variation | P value | Significant |
|---|---|---|
| Interaction | 0.6274 | ns |
| **IND** | **<0.0001** | **** |
| Sex | 0.3119 | ns |

**C. Tar Heel**

| Variation | P value | Significant |
|---|---|---|
| Interaction | 0.5492 | ns |
| **IND** | **<0.0001** | **** |
| Sex | 0.54334 | ns |

**C. Charles River**

| Variation | P value | Significant |
|---|---|---|
| Interaction | 0.4343 | ns |
| **IND** | **<0.0001** | **** |
| Sex | 0.4280 | ns |

**D. Tar Heel**

| Variation | P value | Significant |
|---|---|---|
| Interaction | 0.0790 | ns |
| **IND** | **0.0023** | ** |
| Sex | 0.0790 | ns |

**D. Charles River**

| Variation | P value | Significant |
|---|---|---|
| Interaction | 0.2210 | ns |
| IND | 0.0550 | ns |
| Sex | 0.2210 | ns |

**E. Tar Heel**

| Variation | P value | Significant |
|---|---|---|
| Interaction | 0.4955 | ns |
| **IND** | **0.0009** | *** |
| Sex | 0.4955 | ns |

**E. Charles River**

| Variation | P value | Significant |
|---|---|---|
| Interaction | 0.1616 | ns |
| **IND** | **0.0005** | *** |
| Sex | 0.1616 | ns |

**F. Tar Heel**

| Variation | P value | Significant |
|---|---|---|
| Interaction | 0.2381 | ns |
| **IND** | **0.0098** | ** |
| Sex | 0.2247 | ns |

**F. Charles River**

| Variation | P value | Significant |
|---|---|---|
| Interaction | 0.3467 | ns |
| **IND** | **<0.0001** | **** |
| Sex | 0.4384 | ns |

**G. Tar Heel**

| Variation | P value | Significant |
|---|---|---|
| Interaction | 0.7556 | ns |
| **IND** | **0.0384** | * |
| Sex | 0.7446 | ns |

**G. Charles River**

| Variation | P value | Significant |
|---|---|---|
| Interaction | 0.1570 | ns |
| IND | 0.0643 | ns |
| Sex | 0.1577 | ns |

**H. Tar Heel**

| Variation | P value | Significant |
|---|---|---|
| Interaction | 0.2715 | ns |
| **IND** | **0.0008** | *** |
| Sex | 0.2591 | ns |

**H. Charles River**

| Variation | P value | Significant |
|---|---|---|
| Interaction | 0.1999 | ns |
| **IND** | **0.0105** | * |
| Sex | 0.2313 | ns |

**Fig S1. Indomethacin-induced ileal toxicity in in-house bred Tar Heel and commercial Charles River mice. A**. Scheme of experimental design in which in-house bred "Tar Heel" and commercially purchased "Charles River" C57BL/6 male and female mice were treated with indomethacin (IND; 10 mg/kg body weight) or vehicle for 3 days, and then euthanized 24 h after the last dose (dark arrow). Fecal material were collected at day 0, as indicated. **B.** Percent of day 0 body weight values show significant weight loss in IND treated male (M) and female (F) Tar Heel and Charles River mice at day 3. Additionally, a significant interaction between IND treatment and animal sex was observed in Tar Heel mice. **C.** Fecal lipocalin-2 (LCN2) levels at day 3 are significantly higher in IND treated male (M) and female (F) Tar Heel and Charles River mice. **D.** Ulcer numbers in the ileum are significantly higher in IND treated male Tar Heel mice but not in the other treatment groups. **E.** Inflammation scores in the ileum were significantly higher in IND treated male Tar Heel mice and in IND treated male and female Charles River mice. **F-H.** Gene expression of pro-inflammatory cytokines *Tnf-α, Il-6,* and *Il-1β* normalized by *β-actin* in the ileum are significantly higher in IND treated Tar Heel and Charles River mice of both sexes compared to vehicle treated animals. Analysis of GI toxicity in male and female mouse experiments was performed by two-way ANOVA according to sex type and drug (indomethacin), followed by Tukey-Kramer's method. For the comparison between two treatment groups, statistical significance was determined using two-side t-test. *P<0.05, **P<0.01, ***P<0.001, ****P<0.0001.

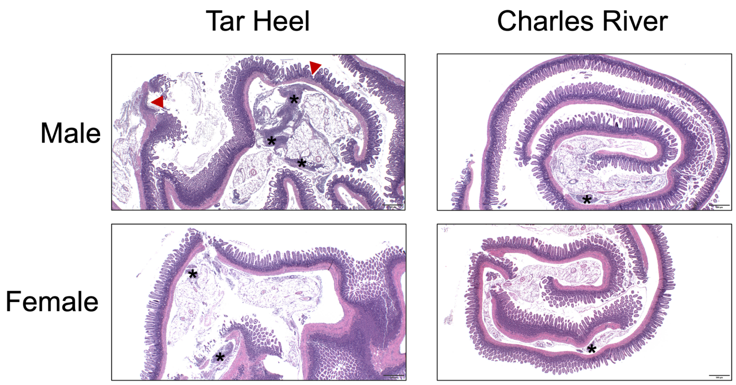


**Fig. S2. Indomethacin-induced ulceration and inflammation in ileum.** Representative H&E histological images from male and female Tar Heel and Charles River mice with ulcers indicated by red triangles and inflammation indicated with asterisks (scale bar = 500 μm).
